# Blood neurofilament light chain levels are associated with disease progression in a transgenic SCA3 mouse model

**DOI:** 10.1101/2023.02.28.530463

**Authors:** David Mengel, Isabel G. Wellik, Kristen H. Schuster, Sabrina I. Jarrah, Madeleine Wacker, Naila S. Ashraf, Gülin Öz, Matthis Synofzik, Maria do Carmo Costa, Hayley S. McLoughlin

## Abstract

Increased neurofilament light (NfL) protein in biofluids is reflective of neurodegeneration and has gained interest as a biomarker across neurodegenerative diseases. In spinocerebellar ataxia type 3 (SCA3), the most common dominantly inherited ataxia, patients exhibit progressive NfL increases in peripheral blood when becoming symptomatic, remaining stably elevated throughout further disease course. However, progressive NfL changes are not yet validated in relevant preclinical SCA3 animal models, hindering its application as a biomarker during therapeutic development. We used ultra-sensitive single-molecule array (Simoa) to measure blood NfL over disease progression in the YACQ84 mouse, assessing relationships with measures of disease severity including age, CAG repeat size, and magnetic resonance spectroscopy. We show that YACQ84 mice exhibit increased blood NfL, concomitant with ataxia-related motor deficits and correlated with neurometabolite abnormalities. Our findings establish natural history progression of NfL increases in the preclinical YACQ84 mouse, further supporting the utility of blood NfL as a peripheral neurodegeneration biomarker and informing coinciding timelines of different measures of SCA3 pathogenesis.

**Summary statement:** Peripheral blood of SCA3 YACQ84 mice exhibits increased abundance of neuronal-specific NfL protein directly associating with disease progression, providing an accessible disease biofluid biomarker to interrogate in preclinical therapeutic studies.

## Introduction

Neurofilament light chain (NfL) is the most abundant subunit of neurofilaments, which are neuron-specific cytoskeletal proteins with crucial roles in structural integrity, electrical signal conduction, and synaptic function of neurons (Gaetani et al., 2019). Of the heavy, medium, and light chains that compose neurofilaments, NfL is the most soluble and can be actively or passively released by cells and detected in biofluids including the cerebrospinal fluid (CSF) (Wilke et al., 2020b, Li et al., 2019, Peng et al., 2022, Garcia-Moreno et al., 2022, Jansen-West et al., 2022, Johnson et al., 2021) and both peripheral blood plasma (Wilke et al., 2020b, Johnson et al., 2018, Peng et al., 2022, Garcia-Moreno et al., 2022, Jansen-West et al., 2022, Coarelli et al., 2021, Byrne et al., 2017) and serum (Wilke et al., 2018, Wilke et al., 2020b, Li et al., 2019, Peng et al., 2022, Soylu-Kucharz et al., 2017). NfL has been shown to be constantly released from neurons to CSF and blood biofluids at low levels in an age-dependent manner, with increased release associated with aging in healthy individuals (Gaetani et al., 2019, Khalil et al., 2020). Due to a key role in providing structural support during radial axon growth, NfL is especially highly expressed in large, myelinated axons and is additionally released at high levels in response to axonal damage (Peng et al., 2022, Gaetani et al., 2019). Elevated levels of NfL in biofluids are considered a strong biomarker for the rate of neuronal turnover, e.g. due to death and degeneration (Benatar et al., 2022, Wilke et al., 2022b, Wilke et al., 2022a, Wilke et al., 2020b) therefore, NfL has been investigated as a biomarker in individuals afflicted with neurodegenerative diseases.

Increased biofluid NfL has been observed in patients with Alzheimer’s disease (Molinuevo et al., 2018, Benedet et al., 2020, Preische et al., 2019), Parkinson’s disease (Mattsson et al., 2019, Hall et al., 2012, Wilke et al., 2020a), parkinsonian disorders (Hall et al., 2012, Hansson et al., 2017), multiple system atrophy-cerebellar subtype (Wilke et al., 2018), sporadic adult-onset ataxia (Wilke et al., 2018), Friedreich’s ataxia (Peng et al., 2022), amyotrophic lateral sclerosis (Weydt et al., 2016, Gaiani et al., 2017), multiple sclerosis (Kuhle et al., 2019, Disanto et al., 2017), Charcot-Maire-Tooth disease (Sandelius et al., 2018), frontotemporal dementia (Rohrer et al., 2016, Wilke et al., 2022b), and Ataxia Telangiectasia (Peng et al., 2022), establishing NfL as a broad biomarker of neurodegeneration. Additionally, there has been increasing evidence of altered NfL in trinucleotide repeat expansion disorders, notably the polyglutamine (polyQ) diseases. These neurodegenerative diseases differ in the implicated gene but share a common mechanism of protein gain of toxic function caused by a polyQ-expansion from a CAG repeat in the disease gene. Of the polyQ diseases, biofluid NfL has been studied in patients with Huntington’s Disease (Johnson et al., 2018, Johnson et al., 2021, Byrne et al., 2017) and multiple spinocerebellar ataxias (SCAs), including SCA1 (Wilke et al., 2018, Coarelli et al., 2021, Peng et al., 2022, Wilke et al., 2022a), SCA2 (Peng et al., 2022, Coarelli et al., 2021), SCA3 (Wilke et al., 2020b, Peng et al., 2022, Garcia-Moreno et al., 2022, Coarelli et al., 2021, Prudencio et al., 2020, Li et al., 2019, Wilke et al., 2018), SCA6 (Peng et al., 2022, Wilke et al., 2018), and SCA7 (Peng et al., 2022, Coarelli et al., 2021).

SCA3, caused by an expanded CAG repeat in the *ATXN3* gene (Kawaguchi et al., 1994), is one of the most common dominantly inherited ataxias worldwide (Gaspar et al., 2001, Schols et al., 2004) and is characterized by cerebellar degeneration and progressive ataxia. With recent advances in therapeutic development for treatment of SCA3 (Costa, 2020, Neves-Carvalho et al., 2020), there has been increased interest in responsive biomarkers informative of disease progression and efficacy of therapeutic intervention. In particular, NfL has been evidenced to hold notable value as a biomarker for SCA3. Patients with SCA3 exhibit significantly higher levels of NfL in serum, plasma, and CSF compared to healthy controls (Wilke et al., 2020b, Li et al., 2019, Peng et al., 2022, Garcia-Moreno et al., 2022, Coarelli et al., 2021, Prudencio et al., 2020, Wilke et al., 2018). Remarkably, SCA3 carriers in pre-ataxic stages also show increased abundance of NfL in biofluids with proximity to age of onset of clinical ataxia presentation (Wilke et al., 2020b). Furthermore, biofluid NfL levels in patients with SCA3 have been found to correlate with various measures of disease progression and severity, including age (Wilke et al., 2020b, Li et al., 2019, Coarelli et al., 2021), CAG repeat size (Wilke et al., 2020b, Coarelli et al., 2021), Scale for Assessment and Rating of Ataxia (SARA) and International Cooperating Ataxia Rating Scale (ICARS) scores (Wilke et al., 2020b, Li et al., 2019, Peng et al., 2022, Garcia-Moreno et al., 2022, Coarelli et al., 2021), and volume of affected brain regions (Li et al., 2019, Peng et al., 2022, Coarelli et al., 2021).

Validation of proposed biomarkers in patients as well as in animal models is imperative to the preclinical assessment of therapeutic efficacy for a specific disease. As such, levels of biofluid NfL have additionally been studied in the context of animal models of neurodegenerative diseases. Mouse models of neurodegenerative diseases, including Huntington’s disease (Soylu-Kucharz et al., 2017) and SCA3 (Wilke et al., 2020b, Garcia-Moreno et al., 2022, Jansen-West et al., 2022, Haas et al., 2022, Lin et al., 2022, Costa et al., 2020), have been shown to exhibit NfL changes similar to those seen in patients. Although levels of NfL are known to change in an age-dependent manner in patients, prior to this study, the association of blood NfL levels with molecular markers of cerebellar pathology and ataxia-like behavior abnormalities in a transgenic mouse model of SCA3 have not yet been investigated. The YACQ84 is the most frequently used preclinical mouse model of SCA3, as it expresses the full-length human mutant *ATXN3* gene harboring a polyQ-encoding CAG repeat of about 84 trinucleotides and exhibits ataxic-like motor deficits and neuropathological signs resembling the human disease (Cemal et al., 2002, Costa et al., 2013, McLoughlin et al., 2018, Moore et al., 2017). In this study, we sought to assess the utility of NfL as a biomarker of disease progression and cerebellar pathology in a preclinical mouse model of SCA3 by characterizing changes in blood NfL levels throughout disease progression of YACQ84 mice.

## Methods and Materials

### Mouse model and sample collection

All animal procedures were approved by the University of Michigan Institutional Animal Care and Use Committee and conducted in accordance with the United States Public Health Service’s Policy on Humane Care and Use of Laboratory Animals. This study used homozygous YACMJD84.2-C57BL/6 transgenic mice (Q84/Q84), hemizygous YACMJD84.2-C57BL/6 transgenic mice (Q84/WT), and non-transgenic wild type (WT/WT) sex- and age-matched littermate controls. Animal genotypes were determined from tail biopsy DNA collected prior to weaning, and confirmed post-mortem as previously described (Cemal et al., 2002, Costa et al., 2013). The *ATXN3* CAG-trinucleotide repeat length was determined by gene fragmentation analysis (Laragen Inc., Culver City) with *ATXN3* primers (5’-ACAGCAGCAAAAGCAGCAA-3’ and 5’-CCAAGTGCTCCTGAACTGGT-3’). CAG repeat length was calculated as (peak amplicon fragment size – 66)/ 3. Serum from groups of homozygous Q84/Q84 (N = 12 [6 females/ 6 males], 36–68 weeks of age) and WT/WT littermate mice (N = 4 [2 females/ 2 males], 56-64 weeks of age), with previously reported neurochemical concentrations measured by magnetic resonance spectroscopy (MRS) (Costa et al., 2020), were assessed. For serum collection at euthanasia, blood was collected from mice by cardiac puncture and immediately centrifuged at 13,000 rpm for 3 min at room temperature. The serum (supernatant) was collected in 0.6 mL Multivette 600z tubes (Sarstedt Inc.) and stored at −80°C. To collect plasma, mice were euthanized and blood was collected by heart puncture with heparinized syringe or 3.2% sodium citrate syringe. Blood was centrifuged in 0.8 mL lithium heparin separator tubes (Greiner Bio-One) at 3,000 *g* for 20 minutes at 4°C, or at 4,000 *g* for 10 minutes at 4°C for sodium citrate tubes. Plasma was collected, aliquoted and stored at −80°C until use. Blood samples were collected for n≥5 animals per genotype per timepoint. Mouse blood was processed to serum, heparin, or citrate plasma to allow different down-stream analyses. Collection of all blood types from a single mouse is not feasible due to limited amounts of available blood. In each experimental paradigm and respective figure of our manuscript, only one type of blood was analyzed to avoid confounding matrix effects. We have further established that while absolute levels of NfL differ between serum and certain forms of plasma from mouse and humans, the magnitude of the disease-specific change in SCA3 vs controls is comparable (Mengel et al. 2023, in preparation).

### Motor assessments

Motor activity of a separate cohort of SCA3 mice relative to WT littermates was measured in an activity chamber with a photobeam open-field apparatus (San Diego Instruments, San Diego, CA). Weight was measured prior to motor assessment at each timepoint.

Total locomotor activity (x/y-axis beam breaks) and rearing activity (z-axis beam breaks) were measured during 30-minute trials. Experimenters were blinded to genotype during behavioral assessment.

### Neurofilament quantification

Neurofilament light concentrations were measured in technical duplicate using the ultra-sensitive single-molecule array (Simoa) on the Simoa HD-X analyzer (Quanterix, Lexington, Massachusetts). The NF-light Advantage kit was used according to manufacturer’s instructions (Quanterix). Mouse blood was spun at 10,000 x *g* for 5 minutes at 4°C and diluted 1 in 4 or 1 in 8 with Quanterix NfL sample buffer for subsequent analysis (dilution linearity of the assay for this concentration range in mouse blood has been established by the Synofzik lab, unpublished results). The lower limit of quantitation (LLoQ) of the assay was defined as the lowest standard: (i) with a signal higher than the average signal for the blank plus 9 SDs, and (ii) allows a percent recovery ≥100 ± 20 %. All measurements were above the LLoQ of the assay and their technical replicates produced an %CV of less than 15%. The LLoQ was defined as 0.48 pg/mL. The repeatability of the NfL assay for two internal control blood samples was determined as 3.4% and 7.1%. Experimenters were blinded to genotype and phenotype prior to analysis.

### Statistical methods

Identification of outliers for all datasets was performed using a ROU Q=1%, and outliers were removed from data analysis. Blood NfL levels were compared between mouse genotypes by linear regression, adjusting for age in the group of aged animals or by two-way ANOVA with post-hoc Tukey multiple comparison’s test in the other groups of animals. Association of NfL levels with cerebellar neurochemical concentrations and age was assessed using Spearman’s rank correlations. Weight and motor activity were compared between mouse genotypes by two-way ANOVA mixed-effects analysis with post-hoc Tukey multiple comparison’s test. All statistical analyses were performed using GraphPad Prism, version 9 (LaJolla, CA). The significance threshold was set to a two-sided *p* < 0.05.

## Results

### Aged homozygous YACQ84 transgenic mice show increased levels of blood NfL

Because clinical trials for SCA3 are routinely preceded by preclinical assays and interventions conducted in mouse models of this disease, it is essential that these models show biofluid biomarkers and biomarker changes like patients with SCA3. Hence, we sought to evaluate whether aged SCA3 YACQ84 transgenic mice expressing the full-length human *ATXN3* disease gene (Cemal et al., 2002), frequently used in preclinical trials for SCA3 (Rodriguez-Lebron et al., 2013b, Ashraf et al., 2019, McLoughlin et al., 2018, Moore et al., 2017, Costa et al., 2013), replicate the increased blood NfL levels observed in patients with SCA3 (Wilke et al., 2020b, Li et al., 2019, Wilke et al., 2018, Peng et al., 2022, Garcia-Moreno et al., 2022, Coarelli et al., 2021).

Using established Simoa-based protocols (Wilke et al., 2020b), we measured blood NfL levels and correlated them with cerebellar neurochemical and pathological alterations (Costa et al., 2020). We used blood samples collected at the time of death immediately following MRS scanning from homozygous YACQ84 (Q84/Q84) mice (average age: 52.3 weeks of age) and their non-transgenic wild type (WT/WT) littermates (average age: 60.0 weeks of age). Aged Q84/Q84 mice replicated the findings in SCA3 patients (Wilke et al., 2020b) by showing on average more than double (216.1 ± 23.40 (SEM) pg/mL) the serum NfL levels displayed by WT/WT mice (87.3 ± 26.87 pg/mL) (Fig. 1A). NfL levels in the two groups of mice did not correlate with age (Fig. 1B) or with CAG repeat size of Q84/Q84 mice (Fig. 1C). However, because serum NfL levels are known to increase with age in healthy individuals (Byrne et al., 2017) as well as with age in SCA3 (Wilke et al., 2020b), the comparison of NfL levels between mouse groups was corrected for this co-variable.

**Figure 1.**
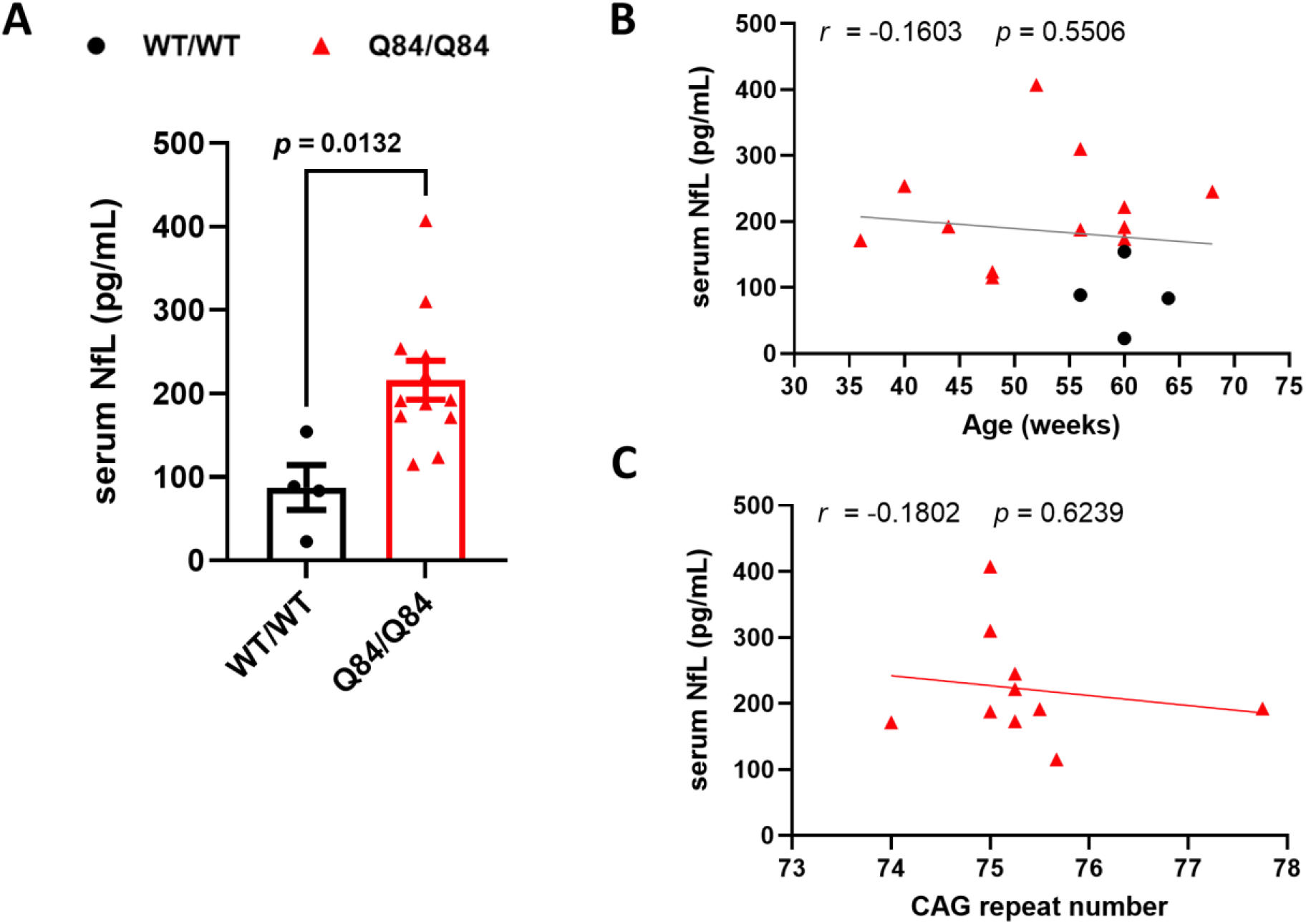
Serum NfL levels are increased in aged Q84/Q84 mice. **(A)** Serum NfL levels of 36–68 weeks-old homozygous Q84/Q84 mice (N=12) are more than double compared to their 50-64 weeks-old WT/WT littermates (N=4). Bars represent mean ± SEM. The displayed *p* value between mouse groups was obtained by linear regression adjusted for age (F=0.0132). Age did not have an effect in the observed differences of NfL abundance between the mouse groups (*p*=0.8860; F=0.02143). **(B)** Serum NfL levels are not correlated with mouse age in the current sample. **(C)** Serum NfL levels of Q84/Q84 mice are not correlated with the CAG repeat size. Associations performed using the Spearman’s rank correlation. *r*, Spearman *r*. Statistical significance was considered for *p* < 0.05.

### Blood NfL levels in homozygous YACQ84 mice correlate with neurochemical signs of cerebellar damage

We have previously reported that decreased levels of N-acetylaspartate (NAA), myo-Inositol (myo-Ins) and total choline (tCho), and increased levels of glutamine (Gln) are potential MRS biomarkers of cerebellar neuronal/axonal injury, demyelination, and astrogliosis in Q84/Q84 mice (Costa et al., 2020). To determine whether increased blood NfL levels in Q84/Q84 mice are associated with cerebellar damage, we correlated individual blood NfL levels with the previously reported neurochemical levels in the cerebellar vermis of the same mice (Costa et al., 2020).

While blood NfL concentration did not correlate with levels of cerebellar total NAA (tNAA) (Fig. 2A) or myo-Ins (Fig. 2B) in aged Q84/Q84 mice, it did correlate inversely with cerebellar abundance of tCho (Fig. 2C), evidenced to be indicative of oligodendrocyte abnormalities (Costa et al., 2020), and directly with cerebellar levels of Gln (Fig. 2D), linked to astrogliosis in prior MRS studies (Öz, 2016), as previously noted. These results suggest that high levels of serum NfL could be driven by the same shared underlying factors as cerebellar axonopathy/demyelination and astrogliosis. Blood NfL levels were not associated with the abundance of other measured neurochemicals (Supplemental Figure 1).

**Figure 2.**
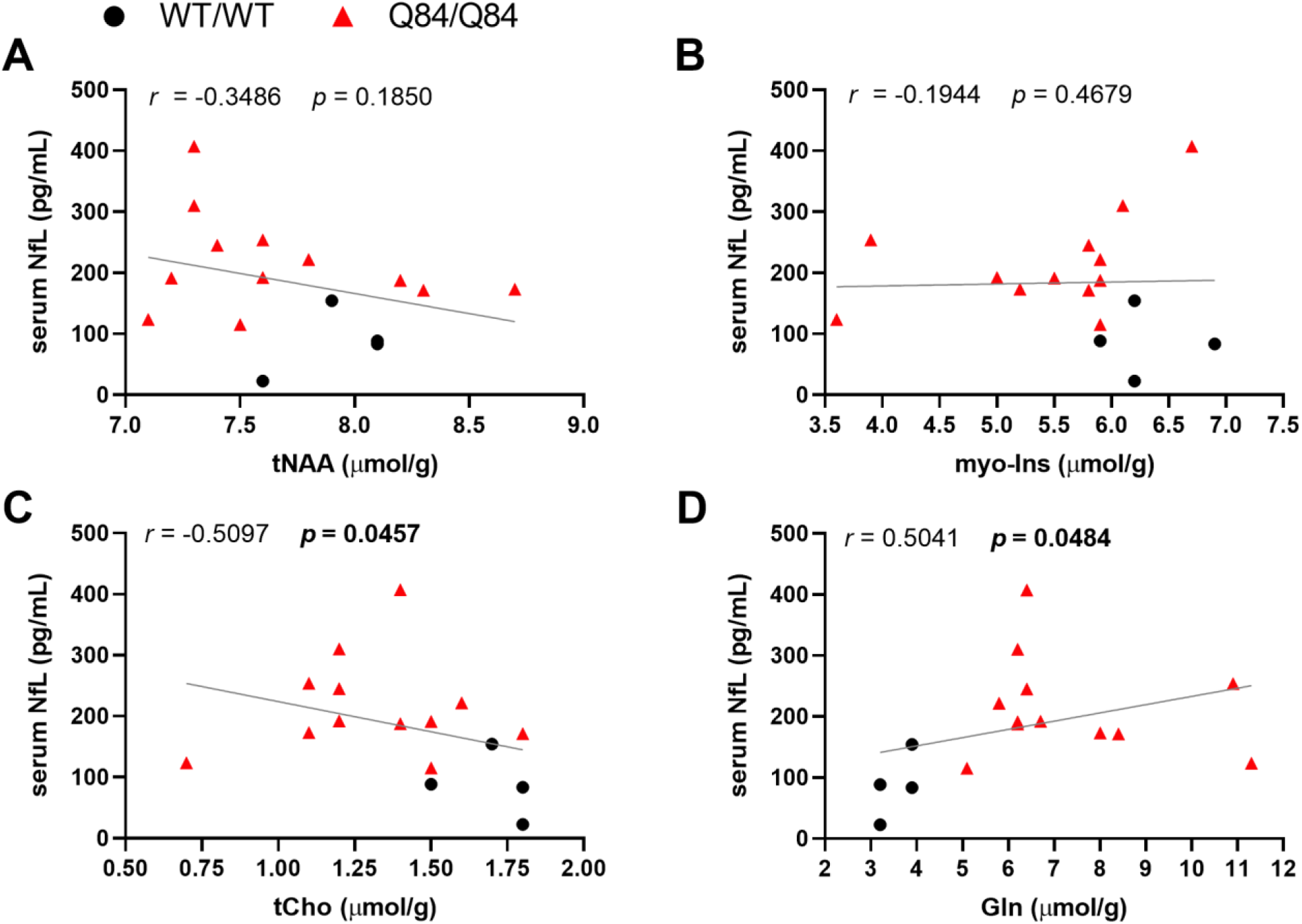
Serum NfL levels are inversely associated with cerebellar total Choline (tCho) and directly associated with cerebellar Glutamine (Gln) levels in Q84/Q84 and WT/WT littermate mice. Scatter plots of serum NfL levels with **(A)** total *N*-acetylaspartate (tNAA), **(B)** myo-Inositol (myo-Ins), **(C)** tCho, and **(D)** Gln. Associations performed using Spearman’s rank correlation. *r*, Spearman *r*. Statistical significance was considered for *p* < 0.05.

### Homozygous YACQ84 mice exhibit ataxia-like motor deficits and weight phenotype at early stages of disease

After observing elevated levels of blood NfL at late-stage disease that also correlated with select neurochemical markers of SCA3 pathology, we aimed to evaluate the progression of motor behavioral deficits and the NfL levels during the disease course. Symptomatic patients with SCA3 are known to experience progressive loss of motor control with disease progression, a behavioral phenotype that is recapitulated by SCA3 mouse models (Cemal et al., 2002, Costa et al., 2013, McLoughlin et al., 2018, Moore et al., 2017). To study the temporal trajectory of ataxia-like motor deficits in our mouse model, we assessed total locomotor activity and rearing activity (open-field testing), as well as weight (Fig. 3A-C) in homozygous Q84/Q84, hemizygous Q84/WT and WT/WT mice at 3, 8, and 14 weeks of age. At 3 weeks of age, locomotor and rearing activities and weight were not significantly different between genotypes (Fig. 3). On the other hand, homozygous Q84/Q84 mice exhibited significantly lower weight compared to WT/WT mice at both 8 and 14 weeks of age, aligning with previous characterization of a weight phenotype in SCA3 mice (Cemal et al., 2002, Costa et al., 2013, Schuster et al., 2023) (Fig. 3A). Consistent with previous findings of symptomatic onset of motor deficits in open field assessment at 4 week of age in homozygous Q84/Q84 mice (Schuster et al., 2023), homozygous Q84/Q84 mice exhibited significant motor deficits compared to WT/WT mice at both 8 and 14 weeks of age, as indicated by decreased total locomotor (Fig. 3B) and rearing activity (Fig. 3C). Hemizygous Q84/WT mice exhibited an intermediate weight and motor phenotype compared to WT/WT and homozygous Q84/Q84 mice (Fig. 3A-C).

**Figure 3.**
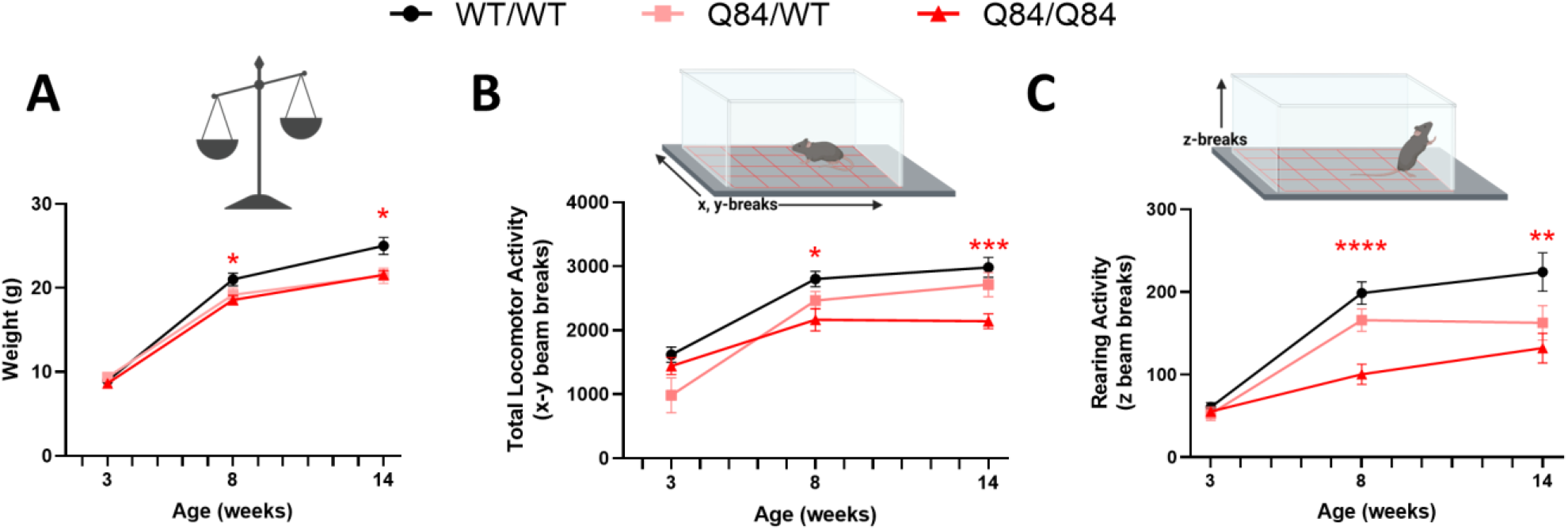
YACQ84 mice exhibit a progressive behavioral phenotype including motor deficits. **(A)** Weight and **(B,C)** behavioral phenotypes were assessed in a separate cohort of mice at 3, 8, and 14 weeks of age, with n≥12 mice for each timepoint. Plots represent mean ± SEM, with behavioral and weight measurements compared between groups by a two-way ANOVA mixed effects analysis with a post-hoc Tukey multiple comparisons test. Statistical significance was considered for *p* < 0.05 (\**p* < 0.05; \*\**p* < 0.01; \*\*\**p* < 0.001, ****p<0.0001). **(B)** Total locomotor activity assessed by x-y beam breaks in activity chamber. **(C)** Rearing activity assessed by z beam breaks in activity chamber.

### YACQ84 mice show age- and disease gene dosage dependent increases in blood NfL, which is coincident with the onset of ataxia-like motor deficits

After having established the onset of ataxia-like motor deficits at an age of 8 weeks, we asked whether temporal trajectories of peripheral NfL levels were correlated to the occurrence of behavioral deficits. To this end, we measured blood NfL levels throughout disease progression at 3, 8, 16, and 45 weeks of age in a separate cohort of mice (Fig. 4A). At 3 weeks of age, there was no significant difference in NfL concentration between homozygous Q84/Q84, hemizygous Q84/WT, and WT/WT mice (Fig. 4B). Beginning at 8 weeks of age, Q84/Q84 mice exhibited significantly increased levels of blood NfL compared to Q84/WT and WT/WT mice (Fig. 4B). At both 8 and 16 weeks, homozygous Q84/Q84 mice had significantly higher plasma NfL levels than hemizygous Q84/WT and WT/WT mice (Fig. 4B). Notably, the timeline of fully penetrant motor deficits in homozygous Q84/Q84 SCA3 mice aligns with this prominent disease feature of a significant NfL increase at 8 weeks in Q84/Q84 mice. While our data demonstrates coinciding timelines of motor deficits and blood NfL increases, future studies will need to establish direct correlation in synonymous cohorts.

**Figure 4.**
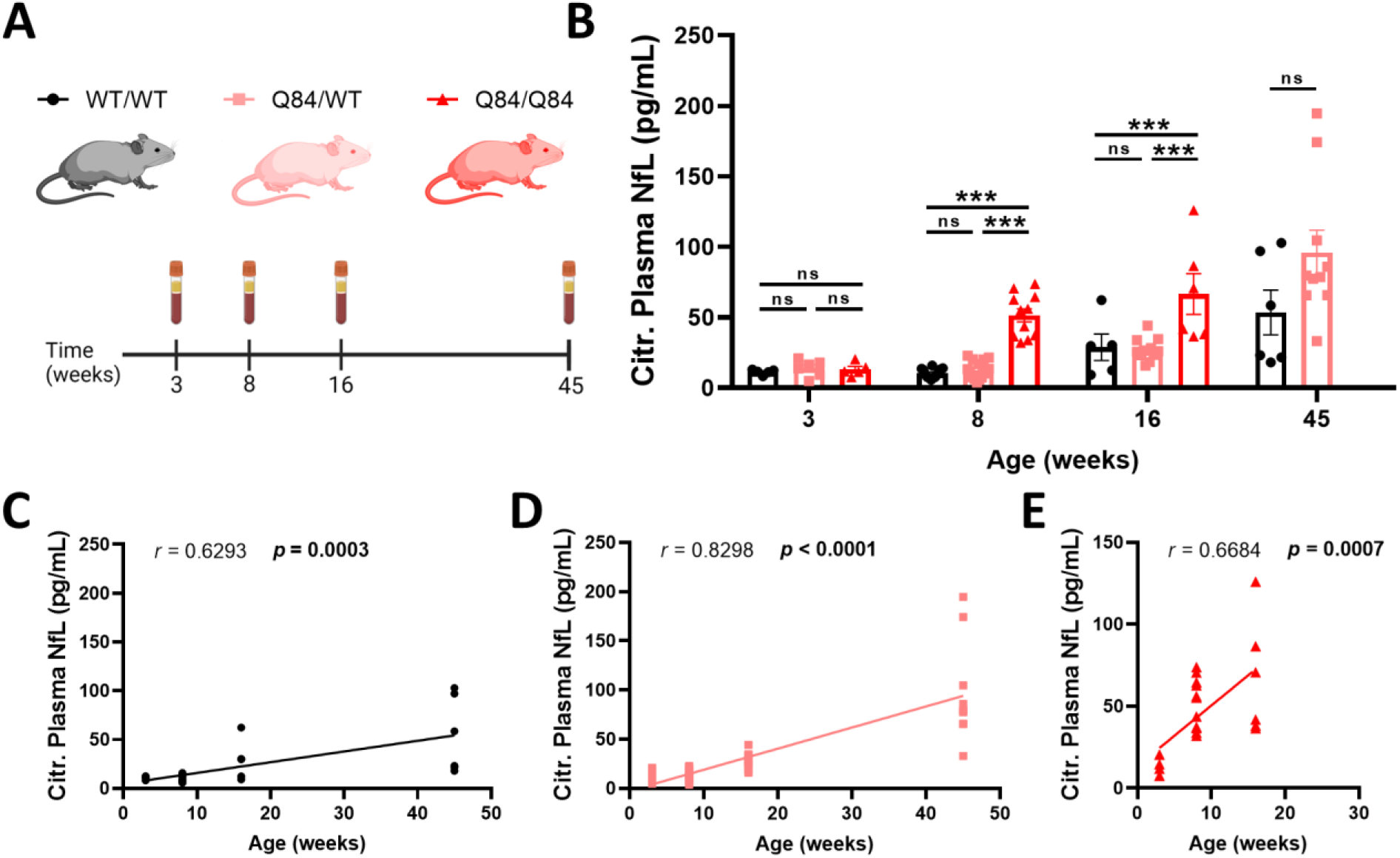
Age and SCA3-dose dependent plasma NfL increase in YACQ84 mice. **(A)** Cross-sectional blood samples from wildtype, hemizygous, and homozygous SCA3 mice were collected at 3, 8, 16, and 45 weeks of age followed by plasma isolation. Blood was collected for n≥5 mice per group at each timepoint, except for homozygous SCA3 mice at 45 weeks. **(B)** NfL concentration (pg/mL) in citrate-treated plasma samples was determined by Simoa. Data are represented as mean ± SEM. Graphs represent statistical data comparison between groups, as assessed by a two-way ANOVA with post-hoc Tukey multiple comparisons test. Statistical significance was considered for *p* < 0.05 (*p<0.05; **p<0.01; ***p<0.001). Correlation between plasma NfL concentration and age in **(C)** WT/WT, **(D)** hemizygous Q84/WT, and **(E)** homozygous Q84/Q84 mice was assessed by Spearman’s rank correlation. *r*, Spearman *r*. Statistical significance was considered for *p* < 0.05.

Following our cross-sectional analysis, we checked if blood NfL levels also directly correlate with mouse age. Unsurprisingly, we found a linear correlation of blood plasma NfL levels with age in WT/WT (Fig. 4C), Q84/WT (Fig. 4D), and Q84/Q84 mice (Fig. 4E), demonstrating a progressive age-dependent increase in NfL concentration over time, as previously demonstrated in patients with SCA3 as well as healthy individuals. Similar to aged Q84/Q84 mice (Fig. 1C), blood NfL levels did not correlate with CAG repeat size in Q84/Q84 mice (Supplemental Fig. 2), thus plasma NfL levels in Q84/Q84 mice were not corrected for CAG repeat size variance. These data indicate that previously established progressive motor impairments track with blood NfL increases over the course of disease, thus aligning molecular and behavioral evidence of progressive neurodegeneration in the Q84 SCA3 mouse model.

## Discussion

Therapeutics for treatment of currently incurable progressive neurodegenerative diseases, including SCA3, have recently advanced to preclinical (Ashraf et al., 2019, McLoughlin et al., 2018, Nobrega et al., 2013, Costa et al., 2013, Rodriguez-Lebron et al., 2013a, Kourkouta et al., 2019, Toonen et al., 2017) and clinical trials (ClinicalTrials.gov Identifiers: NCT05160558, NCT03701399, NCT05490563, NCT04399265), leading to a more immediate imperative for validation of biomarkers of disease progression that can be used to asses therapeutic efficacy. However, it is first crucial to provide validation of such neurodegenerative biomarkers that span both patients and relevant preclinical model systems. Therefore, we used the YACQ84 mouse, known to exhibit clinically relevant aspects of SCA3 pathogenesis (Cemal et al., 2002, Costa et al., 2013, McLoughlin et al., 2018, Moore et al., 2017), to investigate blood NfL changes throughout disease progression. The present study demonstrates that YACQ84 mice recapitulate trends in blood NfL in patients with SCA3, specifically that aged homozygous YACQ84 transgenic mice show increased levels of blood NfL which correlate with neurochemical signs of cerebellar damage. We additionally describe age and gene dose-dependent increases in blood NfL throughout disease progression, aligned with onset of motor impairments. These findings provide informative natural history of NfL biofluid changes in the YACQ84 mouse, validating the neurodegenerative biomarkers in this frequently used SCA3 preclinical model and advising increased opportunity for future preclinical trials in other neurodegenerative mouse models.

Studies in patients with SCA3 have demonstrated trends in NfL increases between the pre-ataxic stage to symptomatic ataxia onset (Wilke et al., 2020b, Li et al., 2019, Peng et al., 2022), and stably increased levels during the symptomatic ataxic phase. These trends are reflective of increased neuronal turnover during the pre-ataxic phase and stably increased neuronal turnover during the symptomatic ataxic phase (Wilke et al., 2020b). Our findings in aged homozygous Q84/Q84 mice reveal synonymous trends, with significantly increased blood NfL levels at late-stage disease. Interestingly, no correlation was found between age and serum NfL in aged homozygous Q84/Q84 mice. However, studies of NfL changes in patients with SCA3 have revealed decreased magnitude of NfL increase over time (Li et al., 2019, Peng et al., 2022, Wilke et al., 2020b), suggesting a plateau effect, i.e. with patients with SCA3 exhibiting stably elevated NfL compared to healthy individuals at late age, yet no longer exhibiting a further acceleration of neuronal turnover over time. Thus, the lack of association between age and biofluid NfL in these mice may be explained by a NfL level plateau at late-stage disease.

Cerebellar neurometabolite abnormalities assessed by MRS, directly prior to biofluid sample collection, correlated with NfL in aged Q84/Q84 mice. In particular, increased serum NfL correlated with decreases in total choline and increases in glutamine, respectively suggestive of oligodendrocyte impairments and gliosis. Thus, the observed correlation between serum NfL and cerebellar neurochemicals in aged homozygous Q84/Q84 mice suggests an association between the magnitude of cellular disturbances and blood NfL. Studies in patients with SCA3 have revealed notable correlations between increased serum NfL levels and neurometabolite abnormalities, specifically decreased cerebellar NAA/Cho and NAA/Cr ratios (Chen et al., 2023). Interestingly, we observed no correlation between NfL and tNAA or myo-Ins, markers of neuronal health and glial contributions, respectively. MRS studies in patients with SCA3 (Miranda et al., 2022, Adanyeguh et al., 2015, Chandrasekaran et al.) and SCA3 mice (Miranda et al., 2022) have revealed significant increases in myo-Ins, however these were specifically observed during early-stage disease in mice, in contrast to our aged Q84 mice. Decreased tNAA has been demonstrated in multiple neurodegenerative diseases (Adanyeguh et al., 2015) including SCA3 (Chandrasekaran et al., Miranda et al., 2022) and is classified as a marker of neuronal and axonal degeneration. While the cerebellum is a heavily impacted region in SCA3, studies in patients and mice demonstrate additional degeneration of the brainstem, substantia nigra, spinal cord, and thalamus (Rub et al., 2013), which we did not assess via MRS in this study. Thus, it is plausible that alternative regional sources of neurodegeneration are contributing to serum NfL increases, explaining the lack of correlation between serum NfL and cerebellar tNAA and myo-Ins neurochemicals. In addition, the lack of correlation between cerebellar tNAA and blood NfL might, in part, also be due to the fact that both biomarkers reflect different aspects of neuronal degeneration: while tNAA levels reflect the cellular dysfunction and dendritic atrophy resulting from neurodegeneration, NfL reflects the *rate* of neurodegeneration (Wilke et al., 2022a, Benatar et al., 2022).

Progressive age-dependent increases in biofluid NfL have been observed in patients with SCA3, with large increases prior to symptomatic ataxia onset (Wilke et al., 2020b, Li et al., 2019, Peng et al., 2022). Notably, healthy controls in studies have also revealed progressive increases in NfL with age (Khalil et al., 2020), though to a lesser extent than seen in patients with neurodegenerative diseases (Gaetani et al., 2019). Our results reveal YACQ84 mice to exhibit progressive increases in blood NfL in an age-dependent manner, mirroring trends seen in patients. While WT/WT mice exhibited NfL increases over time, NfL increases in YACQ84 mice were accentuated in a gene dose-dependent manner with homozygous Q84/Q84 mice showing significantly elevated NfL compared to heterozygous Q84/WT mice. Our previous work (Wilke et al., 2020b) has shown similar trends in age-dependent increases in biofluid NfL in the alternate 304Q Knock-in mouse model, which expresses 304 CAG repeats in the endogenous mouse *Atxn3* gene. The heterozygous 304Q model is characterized by symptomatic onset at 8 months, with a pre-symptomatic increase in plasma NfL at 6 months (Wilke et al., 2020b). In the present study, we demonstrate these same trends of age-dependent increase of plasma NfL in both heterozygous and homozygous YACQ84 mice, with a shortened timeline defined by earlier ataxic onset. Thus, the trends we observe in the YACQ84 mouse, a human mutant *ATXN3* transgene overexpression model, are validated by synonymous blood NfL increases in the endogenous expression system of the 304Q Knock-in mouse model, corroborating a model in which symptomatic onset is reliant upon mutant protein expression in a dose-dependent manner. Accordingly, the YACQ84 mouse enables increased efficiency of longitudinal assessment of disease progression and therapeutic efficacy, re-enforcing its utility as a preclinical model.

SCA3 is characterized by loss of motor function with disease progression, making motor deficits a foremost clinical outcome and central to relevant preclinical models. The YACQ84 faithfully exhibits onset of motor deficits with disease progression (Cemal et al., 2002, Costa et al., 2013, Schuster et al., 2023), rescuable by treatment with anti-*ATXN3* antisense oligonucleotides (McLoughlin et al., 2018). Our findings show that the onset of NfL distinction between healthy mice and SCA3 diseased mice coincides with onset of the reduced weight phenotype and significant motor impairments as measured by open-field motor activity. While open-field activity has previous been validated as a preclinical readout of motor impairment in SCA3 mice (McLoughlin et al., 2018), future investigation of whether NfL increases are relatable to deficits in additional motor tasks would serve to further inform the timeline of disease progression. As the heterozygous 304Q model is characterized by an increase in plasma NfL at 6 months prior to ataxia symptom onset (Wilke et al., 2020b), a deeper understanding of pre-symptomatic NfL increases in the YACQ84 SCA3 mouse model would establish opportunity for preclinical studies to use NfL as an pre-symptomatic biomarker prior to disease onset.

Cohesively, we provide significant natural history of NfL as a biomarker in the YACQ84 mouse, highlighting onset and progressive increases of NfL and behavioral symptoms over time. In newly characterizing this neurodegenerative disease feature in the YACQ84 mouse, we further validate its use as a preclinical model of SCA3 and establish a basis for future studies to investigate therapeutic timelines that maximize reduction of neurodegeneration in SCA3.

## Acknowledgements

The authors would like to thank Ilya Bezprozvanny for sharing the YACQ84 mice.

## Author Contributions

Conceptualization: DM, GÖ, MS, MCC, HSM

Methodology: DM, MS, MCC, HSM

Formal analysis: DM, IGW, MCC

Investigation: DM, IGW, KHS, SJ, MW, NSA, MCC, HSM

Writing-original draft preparation: IGW, MCC, HSM

Writing-review and editing: DM, IGW, KHS, SJ, MW, NSA, GÖ, MS, MCC, HSM

Funding acquisition: DM, MS, MCC, HSM

---

**Supplemental Figure 1.**
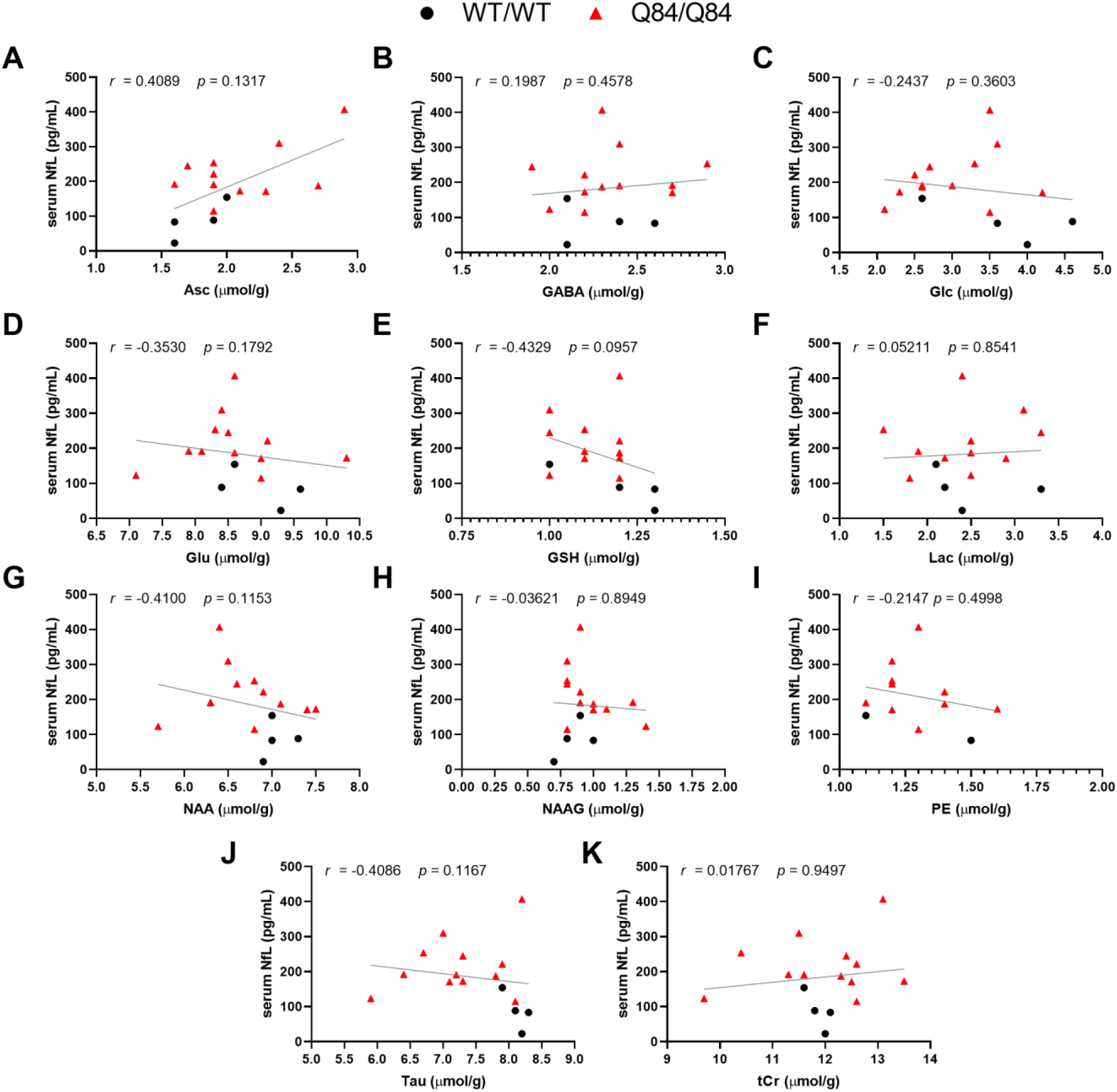
Scatter plots of serum NfL levels not correlated with cerebellar **(A)** Asc, ascorbate; **(B)** GABA, γ-aminobutyric acid; **(C)** Glc, glucose; **(D)** Glu, glutamate; **(E)** GSH, glutathione; **(F)** Lac, lactate; **(G)** NAA, *N*-acetylaspartate; **(H)** NAAG, *N*-acetylapartylglutamate; **(I)** PE, phosphoethanolamine; **(J)** Tau, taurine; **(K)** tCr, total creatine in WT/WT and Q84/Q84 homozygous mice. Line of best fit determined by simple linear regression. Associations performed using Spearman’s rank correlation. *r*, Spearman *r*. Statistical significance was considered for *p* < 0.05.

**Supplemental Figure 2.**
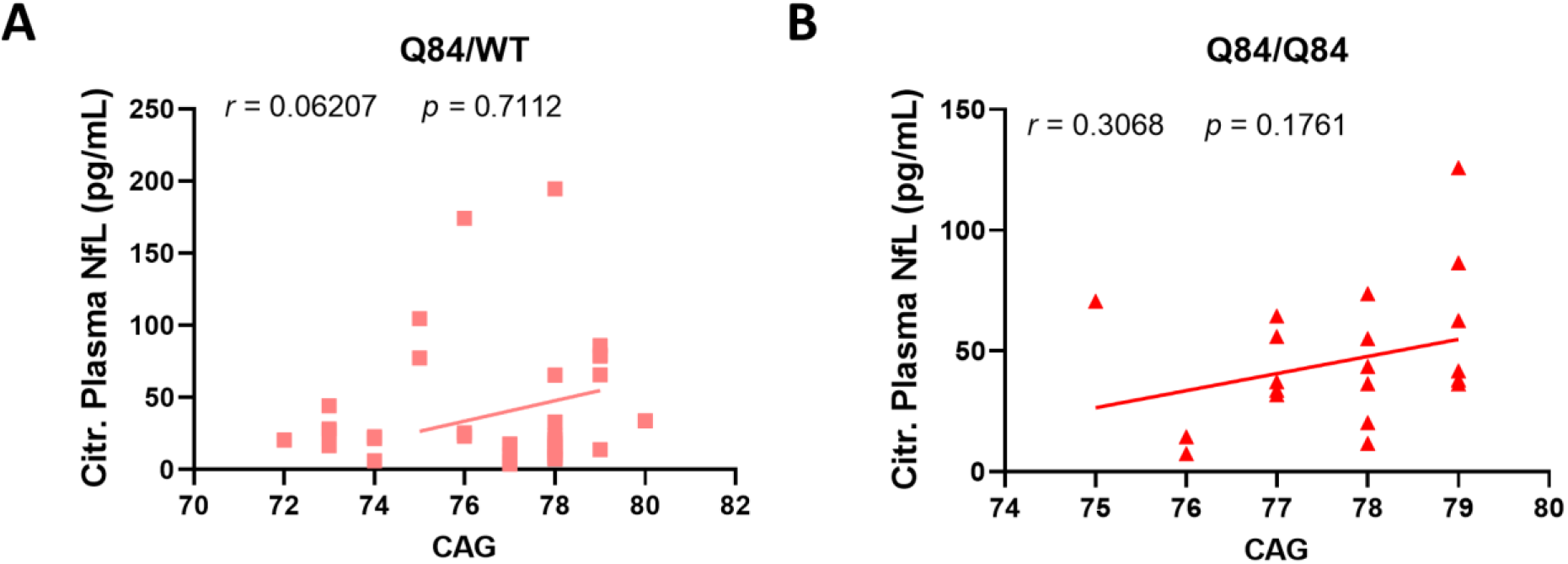
Scatter plots of sodium citrate plasma NfL levels with CAG repeat size in **(A)** Q84/WT hemizygous and **(B)** Q84/Q84 homozygous mice. Associations performed using Spearman’s correlation. *r*, Spearman *r*. Statistical significance was considered for *p* ≤ 0.05.

